## Supplementary Information for "USP22 overexpression fails to augment tumor formation in MMTV-ERBB2 mice but loss of function impacts MMTV promoter activity"

Supporting Information

S1 Fig. Validation of high enrichment of epithelial cells vs fibroblasts in lysates from mammary epithelial cells (MECs) collected from 100-day old NIC mice.

A. Immunoblot analysis of lysates using epithelial (E-cadherin, Cytokeratin 8) and fibroblast (FSP-1) specific markers in 3 biological replicates from each genotype analyzed. NMuMG and NIH3T3 cell lysates were used as positive samples for epithelial cell and fibroblast cell, respectively.

Supporting Materials

S1 Table. qRT-PCR primers

| Gene name | Forward Primer | Reverse Primer |
| --- | --- | --- |
| Human *USP22* | CTCCTGTCTGGTCTGTGAGATG | CAGCAACTTATACGGGATGTGA |
| Human *ERBB2* | TGCTGGACATTGACGAGACAGAGT | AGCTCCCACACAGTCACACCATAA |
| Mouse *Usp22* | GAGAGCAGGATGAATGGGC | CCTTGGTGATAATGGCGTC |
| Mouse/Rat *Erbb2* | TCCCTGCCAGTCCTGAGACC | GTTGTGAGCGATGAGCATGTA |
| Mouse *GR* | GGGCGCCAAGTGATTGCCGCAGT | CCAACCCAGGGCAAATGCCATGA |
| Mouse *PR* | TGGGAAAGTCTTCGACGTGAC | GTGCATCCTTATCCAGGCAGA |
| Mouse *Brg1* | GAGCCAGAACGAGAAGTACCG | CCTCAAGACGAGCAATTTCATCA |
| Human *GAPDH* | CTCCTCCTGTTCGACAGTCAGC | CCCAATACGACCAAATCCGTT |
| Mouse *Pbgd* | GTGTTGCACGATCCTGAAACT | GTTGCCCATCCTTTATCACTGTA |

S2 Table. Antibodies

| Antibody name | Manufacturer | Catalog number |
| --- | --- | --- |
| USP22 | Abcam | ab195289 |
| ERBB2/HER2 | Invitrogen | MA5-13105 |
| FLAG | Sigma | F7425 |
| GAPDH | EMD Millipore | MAB374 |
| Mouse GR | Proteintech | 24050-1-AP |
| Phospho-ERBB2 | Cell Signaling Technology | 2243 |
| Mouse BRG1 | Proteintech | 21634-1-AP |
| ERα | Santa Cruz | sc-542 |
| E-CADHERIN | Thermo Fisher | 13-1900 |
| CYTOKERATIN 8 (CK8) | Developmental Studies Hybridoma Bank (DSHB) | TROMA-1-c |
| FSP-1 | Proteintech | 16105-1-AP |
| SLUG | Proteintech | 12129-1-AP |
| OCCLUDIN | Thermo Scientific | 33-1500 |
| VIMENTIN | Abcam | ab92547 |
| TWIST | Santa Cruz | sc81417 |
| Smooth muscle actin (SMA) | Sigma | A6228 |
| GCN5 | Cell Signaling Technology | 3305 |
| ATXN7L3 | Gift from Dr. László Tora |  |
| ERBB3 | Cell Signaling Technology | 12708 |
| HSP70 | Cell Signaling Technology | 4872 |

| HSP90 | Cell Signaling Technology | 4877 |
| --- | --- | --- |
| HSP90AB1 | Proteintech | 11405-1-AP |
| AKT | Cell Signaling Technology | 4691 |
| Phospho-AKt | Cell Signaling Technology | 4060 |
| ERK1/2 | Cell Signaling Technology | 4695 |
| Phospho-ERK1/2 | Cell Signaling Technology | 4370 |
| H2B | Millipore | 07-371 |
| H2Bub | Cell Signaling Technology | 5546 |
| Rabbit IgG-HRP (from donkey) | Amersham-Cytiva | NA934-1ML |
| Mouse IgG-HRP (from donkey) | Amersham-Cytiva | NA931-1ML |
| Rat IgG-HRP (from goat) | Cell Signaling Technology | 7077 |

S3 Table. USP22 lentiviral shRNA vectors used in the project are from Open biosystems

(currently could be found in Horizon Discovery)

| Name of vector | Cat# | Clone ID |
| --- | --- | --- |
| GIPZ Non-silencing lentiviral shRNA control | RHS4346 |  |
| GIPZ lentiviral human *USP22* shRNA-1 | RHS4430-200200653 | V2LHS_60727 |
| GIPZ lentiviral human *USP22* shRNA-2 | RHS4430-200305445 | V3LHS_377162 |
| GIPZ lentiviral human *USP22* shRNA-3 | RHS4430-200161960 | V2LHS_13647 |
