## Supplementary figures and images for "USP22 overexpression fails to augment tumor formation in MMTV-ERBB2 mice but loss of function impacts MMTV promoter activity"

### Supp Figure 1

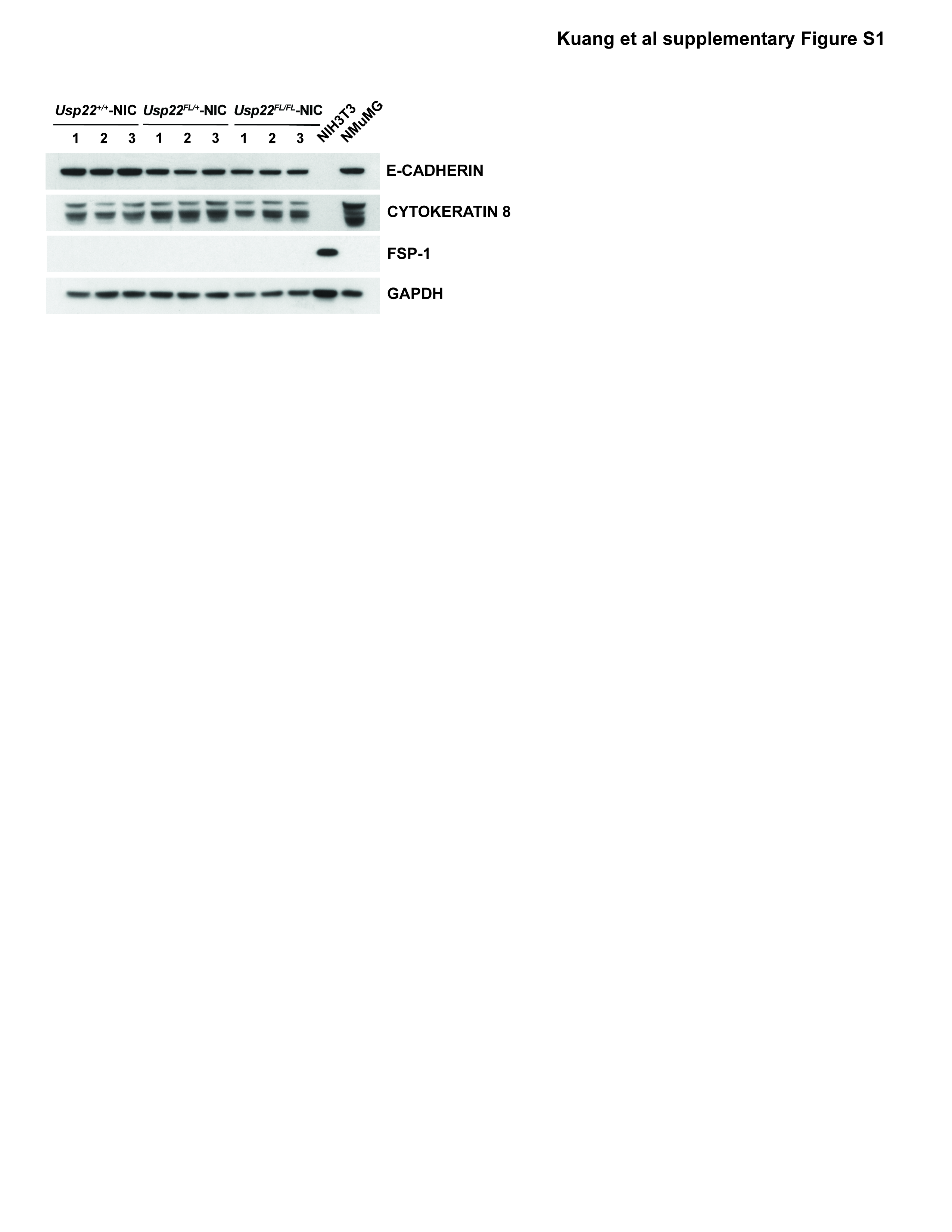
